# Latent inter-organ mechanism of idiopathic pulmonary fibrosis unveiled by a generative computational approach

**DOI:** 10.1101/2023.04.18.537146

**Authors:** Satoshi Kozawa, Kengo Tejima, Shunki Takagi, Masataka Kuroda, Mari Nogami-Itoh, Hideya Kitamura, Takashi Niwa, Takashi Ogura, Yayoi Natsume-Kitatani, Thomas N. Sato

## Abstract

Idiopathic pulmonary fibrosis (IPF) is a chronic and progressive disease characterized by complex lung pathogenesis affecting approximately three million people worldwide. While the molecular and cellular details of the IPF mechanism is emerging, our current understanding is centered around the lung itself. On the other hand, many human diseases are the products of complex multi-organ interactions. Hence, we postulate that a dysfunctional crosstalk of the lung with other organs plays a causative role in the onset, progression and/or complications of IPF. In this study, we employed a generative computational approach to identify such inter-organ mechanism of IPF. The approach works as follows: 1) To find unexpected relatedness of IPF to other diseases of non-lung organs and to identify molecular features that define such relatedness, 2) To identify differentially expressed genes between the lung tissues of IPF vs. those of non-IPF pulmonary disease patients, 3) To detect ligand-receptor relationships across multiple organs and their upstream and downstream signaling pathways in 1) and 2), 4) To generate a map of the inter-organ IPF mechanism with the molecular and cellular resolution. This approach found unexpected molecular relatedness of IPF to neoplasm, diabetes, Alzheimer’s disease, obesity, atherosclerosis, and arteriosclerosis. Furthermore, as a potential mechanism underlying this relatedness, we uncovered a putative molecular crosstalk system across the lung and the liver. In this inter-organ system, a secreted protein, kininogen 1, from hepatocytes in the liver interacts with its receptor, bradykinin receptor B1 in the lung. This ligand-receptor interaction across the liver and the lung leads to the activation of calmodulin pathways in the lung, leading to the activation of interleukin 6 and phosphoenolpyruvate carboxykinase 1 pathway across these organs. Furthermore, we retrospectively identified several pre-clinical and clinical evidence supporting this inter-organ mechanism of IPF. In conclusion, such feedforward and feedback system across the lung and the liver provides a unique opportunity for the development of the treatment and/or diagnosis of IPF. Furthermore, the result illustrates a generative computational framework for machine-mediated synthesis of mechanisms that facilitates and complements the traditional experimental approaches in biomedical sciences.

## Introduction

Idiopathic pulmonary fibrosis (IPF) is a chronic disease characterized by scarring in the interstitium of the lung, affecting 3 – 9 and 4 or less per 100,000 person-years in North America/Europe and South America/East-Asia, respectively^1,2^. Both the incidence and poor prognosis of IPF increase with age^3,4^. Specifically, the median age of the newly diagnosed is 62 years-old and their prognosis is poor – 3 – 5 years of survival rate.

There are two FDA-approved drugs for the treatment of IPF: nintedanib and pirfenidone^1,2^. Nintedanib is a tyrosine kinase inhibitor. Pirfenidone is an inhibitor of TGF-beta production and downstream signaling, collagen synthesis and fibroblast proliferation. Hence, these drugs are regarded as pleiotropic anti-fibrosis drugs. Currently there are no IPF-specific therapeutics. Furthermore, the precise IPF diagnosis requires complex and multiple-types of tests as its overlapping pathologies with other interstitial lung fibrosis diseases^1,2^. These are in part due to the complexity of the IPF pathogenesis and to its ill-defined cellular and molecular mechanisms.

While IPF was classically considered an inflammatory disease, a new picture is emerging^1–6^. The increasing molecular and cellular evidence suggests IPF is driven by an activation of the lung epithelium. In this model, the ectopic activation of the alveolar epithelial cells results in the production of chemokines, growth factors, and extracellular matrix proteins, promoting the migration, growth, and/or differentiation of fibroblasts, and macrophages and other immune cells. Furthermore, various life-style and environmental factors, and also genetic factors are reported to influence the onset, progression, and/or mortality.

While the clinical translation of these recent advancements in understanding the IPF mechanism within the lung tissue is expected, the onset and progression of human diseases involve multiple organs -i.e., the inter-organ mechanism^7–11^. The immune responses occur in a variety of diseases such as metabolic, neoplastic, cardiovascular diseases and also in aging^12–14^. The causative roles of gut microbiota are becoming recognized in an increasing number of diseases^15^. The nervous system influences metabolic states and vice versa^16^. Metabolic dysregulation is a risk factor for many diseases and they also accelerate aging influencing the longevity^17^. Exosomes are another type of systemic factors that are associated with many types of diseases^18^. The interactions of immune cells and lung cells are involved in the pathogenesis of IPF^1,2,4–6^. Furthermore, life-style and aging are critical influencers of IPF^1,5^. Hence, it is conceivable that the inter-organ crosstalk mediated by the immune cells, systemic factors, and/or neural system could be involved in the onset, progression, and/or complications of IPF. However, very little is studied on these possibilities.

Based on this background, we postulate that the cross-talk between the lung and the non-lung organs is a part of the mechanism in the onset and/or progression of IPF. The obvious choice of the approach to test this hypothesis is to examine molecular and cellular changes in non-lung organs that accompany, precede, or follow the pathological changes of the lung in IPF. However, this approach would be difficult as the availability of non-lung tissues from the IPF patients is limited, if any.

The availability of multi-modal omics data of multiple organs and diseases is growing in the public space^19–27^. Such data space, together with computational methods, could allow us to deduce what occurs in the non-lung organs of IPF-patients and to simulate how they are regulated.

Hence, we reasoned that such a large set of multi-modal omics data across many organs and diseases in the human body provides an uncharted biomedical space where an organ-to-organ interaction that causes and/or exacerbates IPF is embedded. To uncover such a latent inter-organ mechanism of IPF, we employ a generative computational approach (also referred to as “generative AI”) – a computational method that can produce various types of contents such as sentences, images, molecular structures, working-hypotheses(models), etc.^28^.

Towards this goal, we designed a generative computational approach as follows:

1. To detect mechanistic relatedness of IPF to non-respiratory/non-pulmonary diseases.
2. To identify molecular features that characterize the relatedness detected in 1.
3. To identify ligand – receptor relationships across multiple organs that are linked to the features identified in 2.
4. To generate a map of the inter-organ mechanism of IPF with molecular and cellular resolution that explains the findings of 1 – 3.

## Results

### Detection of molecular relatedness of IPF to non-respiratory/non-pulmonary diseases

The molecular relatedness of IPF to non-respiratory/non-pulmonary diseases were identified by using the multi-modal generative topic modeling method that we developed and previously reported^29^. The overall design is summarized in Fig. 1, and it works as follows:

**Fig. 1.**
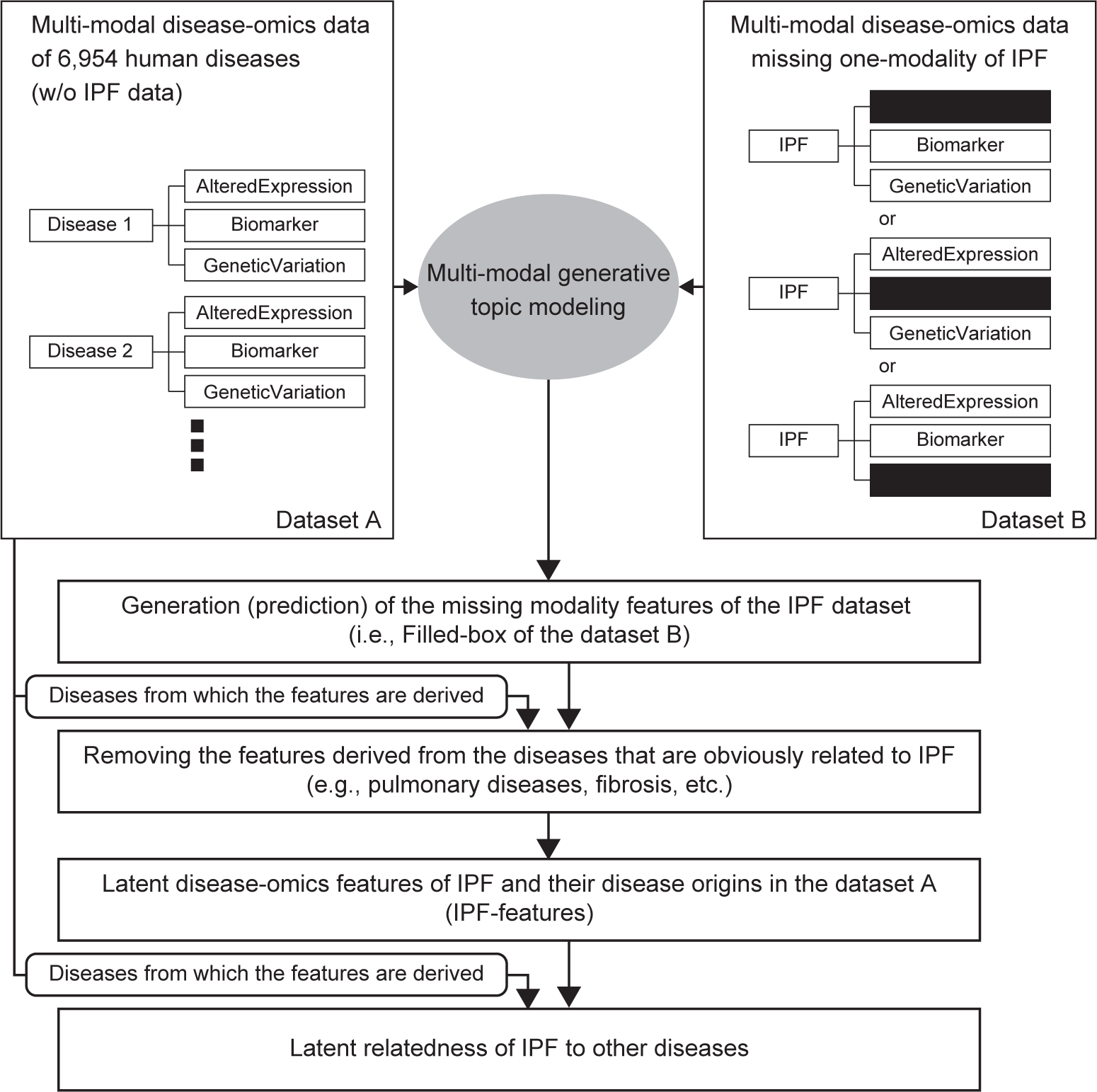
General overview of the multi-modal generative topic modeling approach for IPF. The previously developed method^29^ is adapted to IPF.

Two datasets are used for the multi-modal generative topic modeling: Datasets A and B. Dataset A consists of 6,954 human diseases excluding IPF, each of which is characterized by three disease omics modalities, AlteredExpression (Ae), Biomarker (Bm), and GeneticVariation (Gv), derived from DisGeNET v7.0.^19,27^. “Ae” is the list of genes and proteins of which changes in expressions are associated with a corresponding disease(s). “Bm” is the list of biomarkers which are described for a corresponding disease(s). “Gv” is the list of genes of which mutations are reported for a corresponding disease(s). Dataset B consists of three types of IPF modality combination, each consisting of Bm/Gv (i.e., missing Ae), Gv/Ae (i.e., missing Bm), or Ae/Bm (i.e., missing Gv). The multi-modal generative topic modeling generates (i.e., predicts) the features of the missing modalities. The performance was evaluated by calculating the AUC values as previously described^29^ and they were found to be above 0.8 for all three modalities (Fig. S1). Next, from these computationally generated features, those derived from the modalities of IPF itself and those of obviously IPF-related diseases are removed. The diseases that are obviously related are those of which names contain “Pulmonary”, “Lung”, “Fibrosis”, “Respir**”, “Chest”, “Pneumo**” (** could be any characters). The remaining features are now designated as “latent disease-omics features of IPF (also referred to as IPF-features)”. Moreover, IPF and the diseases from which these IPF-features are derived in Dataset A establish “latent relatedness of IPF to other diseases”.

Using this approach, we identified 83 latent IPF-features (Table S1). The human-organ-expression analysis using THE HUMAN PROTEIN ATLAS v 21.1.^23–25^ (see also Methods section) found that their expression is most enriched in the liver (Fig. 2A, Table S2). Additionally, we also detected the statistically significant (i.e., q-values < 0.05) enrichments in the immune system (bone marrow, lymphoid tissue, blood), the kidney, the thyroid gland, adipose tissue, the prostate, and the placenta. The cellular level analysis found the highest enrichment in the hepatocytes (Fig. 2B, Table S2). In addition, we also detected the statistically significant (i.e., q-values < 0.05) enrichments in Kupffer cells and Hofbauer cells, two types of macrophages found in the liver and the placenta, respectively. These results suggest a possibility that the liver, in particular, hepatocytes and the intra-hepatic immune cells such as Kupffer cells, participate in the pathogenesis of IPF.

**Fig. 2.**
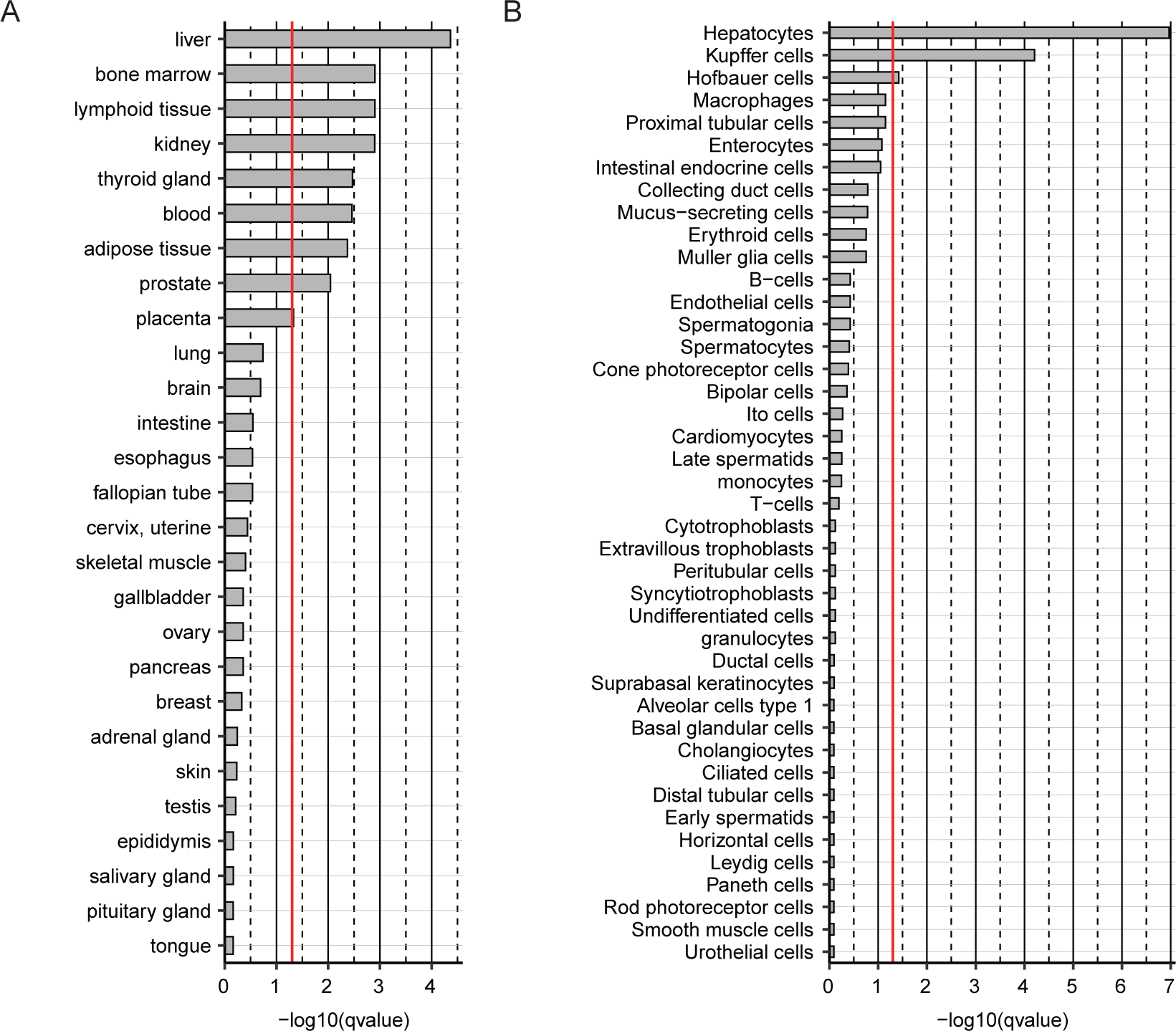
The organ and cell-enrichment analyses of the latent IPF-features. A: The organ enrichment. B: The cell-type enrichment. The enrichment level of the 83 IPF-features in each organ and each cell-type is shown as bar-graph of log10(q-values) in the descending order. The q-value (qvalue)=0.05 (the threshold for the statistical significance) is indicated as a red line in each graph. The raw data are available as Table S2.

### Latent relatedness of IPF to other diseases

Next, we determined the disease-label(s) of the 83 latent IPF-features to identify non-pulmonary/non-respiratory diseases to which IPF is related (Fig. 3, Table S3) (see also Methods section). This analysis found these IPF-features are derived from neoplastic diseases. In addition, they are also labeled with autoimmune disorders, diabetes, Alzheimer’s disease, rheumatoid arthritis, obesity, cardiovascular diseases (atherosclerosis, arteriosclerosis, hypertensive disease, etc.), systemic lupus erythematosus, and multiple sclerosis. The result suggests that these non-pulmonary/non-respiratory diseases are related to IPF at the molecular level.

**Fig. 3.**
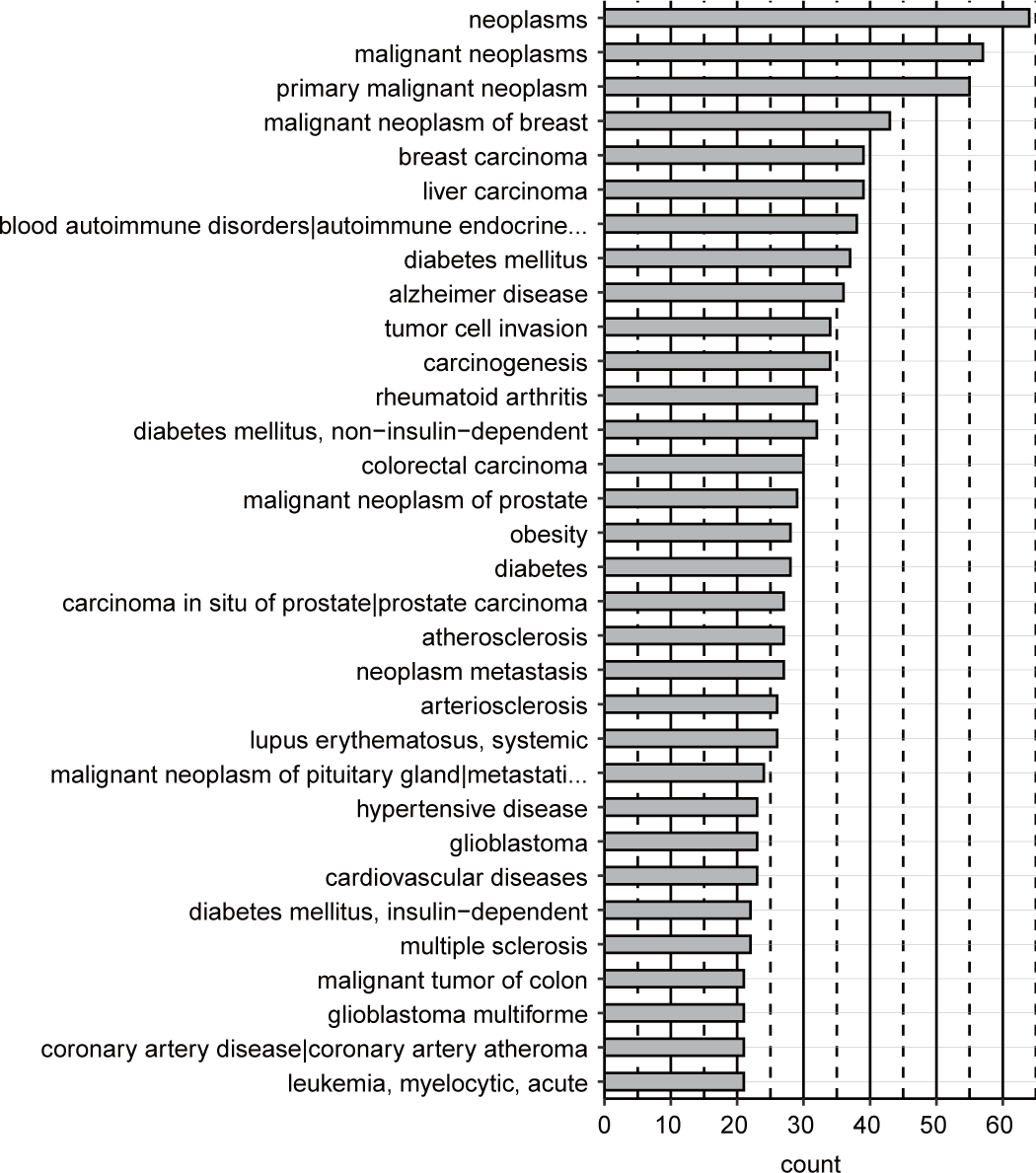
The latent diseases to which IPF is molecularly related. The frequency of the appearance of the 83 IPF-features in each disease is indicated as “count”. Shown are the diseases of which counts are above 20 in the descending order. The long disease names are cut short and indicated as “..” at their ends. The raw data are available as Table S3.

### Inter-organ mechanism of IPF

A putative inter-organ mechanism was computationally generated as described in Fig. 4 and the overall flow is as follows:

1. To identify the ligands in the lung DEgenes (via CellChatDB as above) and to determine the cell-type(s) in the lung where they are expressed.
2. To identify the receptors for the ligands in 1 (via CellChatDB as above) and to determine the non-lung organ(s) and the cell-type(s) where they are expressed.
3. To identify the receptors in the lung DEgenes (via CellChatDB as above) and to determine the cell-type(s) in the lung where they are expressed.
4. To identify the ligands for the receptors in 3 (via CellChatDB as above) and to determine the non-lung organ(s) and the cell-type(s) where they are expressed.
5. To determine the KEGG pathways to which the receptors (identified in 2) and the ligands (identified in 4) belong.
6. To identify the latent IPF-features that belong to the same KEGG pathways as those of 5.
7. To construct the inter-organ map on the basis of 1 – 6 results.

**Fig. 4.**
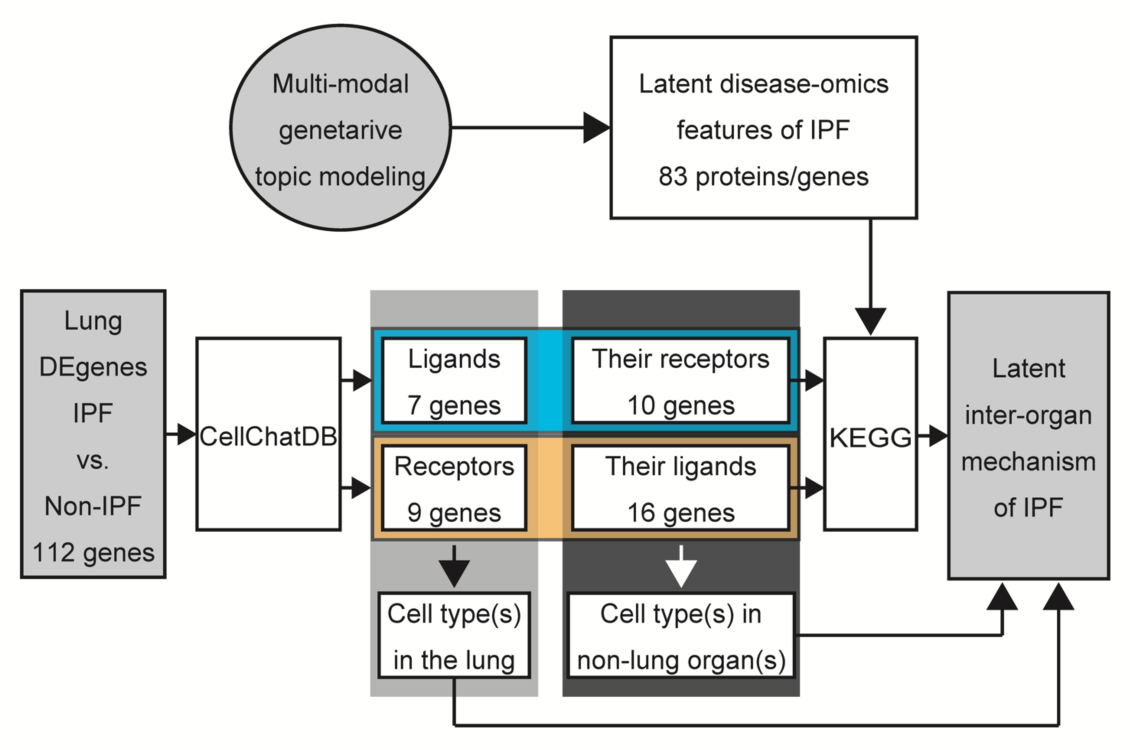
General overview of the computational framework to generate an inter-organ mechanism of IPF. See the Methods section for the detailed step-by-step description. The 83 latent IPF-features and 112 lung DEgenes (IPF vs. non-IPF) are found in Tables S1 and S4, respectively.

The DESeq2 analysis of the lung tissues obtained from 95 IPF and 204 non-IPF lung disease patients (see also Methods section) identified a total of 112 IPF-DEgenes (Table S4). The CellChatDB analysis identified seven ligands (CCL18, CXCL9, CXCL10, CXCL11, IL6, IFNG, SELE) and nine receptors (CXCR3, CXCR5, CXCR6, BDKRB1, CHRNA1, TMIGD3, DSC3, PDCD1, SELE) in the 112 IPF-DEgenes (Tables 1&2). For the seven ligands, there exist eight receptors (ACKR1, CXCR3, ACKR3, IL6R/IL6ST, IFNGR1/IFNGR2, CEACAM1, CD44, GLG1), forming 13 ligand-receptor pairs (CCL18-ACKR1, CXCL9-ACKR1, CXCL10-ACKR1, CXCL11-ACKR1, CXCL9-CXCR3, CXCL10-CXCR3, CXCL11-CXCR3, CXCL11-ACKR3, IL6-IL6R/IL6ST, IFNG-IFNGR1/IFNGR2, SELE-CEACAM1, SELE-CD44, SELE-GLG1) (Table 1). For the nine receptors, there exist 16 ligands (PF4V1, CXCL9, CXCL10, CXCL11, CXCL13, PF4, CXCL16, KNG1, SLURP1, SLURP2, ENTPD1, DSG1, DSG2, CD274, PDCD1LG2, SELPLG), forming 17 ligand-receptor pairs (PF4V1-CXCR3, CXCL9-CXCR3, CXCL10-CXCR3, CXCL11-CXCR3, CXCL13-CXCR3, PF4-CXCR3, CXCL13-CXCR5, CXCL16-CXCR6, KNG1-BDKRB1, SLURP1-CHRNA1, SLURP2-CHRNA1, ENTPD1-TMIGD3, DSG1-DSC3, DSG2-DSC3, CD274-PDCD1, PDCD1LG2-PDCD1, SELPLG-SELE) (Table 2).

**Table 1.** The ligands encoded by the IPF-DEgenes and their receptors

| Ligand (IPF-DEgene) | Receptor | Evidence |
| --- | --- | --- |
| CCL18 | ACKR1 | PMID: 26740381 |
| CXCL9 | ACKR1 | PMID: 26740381 |
| CXCL10 | ACKR1 | PMID: 26740381 |
| CXCL11 | ACKR1 | PMID: 26740381 |
| CXCL9 | CXCR3 | KEGG: hsa04060 |
| CXCL10 | CXCR3 | KEGG: hsa04060 |
| CXCL11 | CXCR3 | KEGG: hsa04060 |
| CXCL11 | ACKR3 | KEGG: hsa04060 |
| IL6 | IL6R / IL6ST | KEGG: hsa04060 |
| IFNG | IFNGR1 / IFNGR2 | KEGG: hsa04060 |
| SELE | CEACAM1 | PMID: 1378450 |
| SELE | CD44 | PMC4571854 |
| SELE | GLG1 | PMID: 11404363 |

The ligands are IPF-DEgenes. The evidence for each ligand-receptor pair is indicated as described in CellChatDB.

**Table 2.** The receptors encoded by the IPF-DEgenes and their ligands

| Ligand | Receptor (IPF-DEgene) | Evidence |
| --- | --- | --- |
| PF4V1 | CXCR3 | KEGG: hsa04060 |
| CXCL9 | CXCR3 | KEGG: hsa04060 |
| CXCL10 | CXCR3 | KEGG: hsa04060 |
| CXCL11 | CXCR3 | KEGG: hsa04060 |
| CXCL13 | CXCR3 | KEGG: hsa04060 |
| PF4 | CXCR3 | KEGG: hsa04060 |
| CXCL13 | CXCR5 | KEGG: hsa04060 |
| CXCL16 | CXCR6 | KEGG: hsa04060 |
| KNG1 | BDKRB1 | KEGG: hsa04080 |
| SLURP1 | CHRNA1 | KEGG: hsa04080 |
| SLURP2 | CHRNA1 | KEGG: hsa04080 |
| ENTPD1 | TMIGD3 | PMID: 21677139 |
| DSG1 | DSC3 | PMID: 27298358 |
| DSG2 | DSC3 | PMID: 27298358 |
| CD274 | PDCD1 | PMID: 23954143 |
| PDCD1LG2 | PDCD1 | PMID: 23954143 |
| SELPLG | SELE | PMC4571854 |

The receptors are IPF-DEgenes. The evidence for each ligand-receptor pair is indicated as described in CellChatDB.

We then searched for the downstream and upstream signaling targets for these ligand-receptor pairs by the KEGG-mining (see Methods section). For the 13 ligand (lung) – receptor (non-lung) pairs, we found six such targets in the latent IPF-features (Table 3) - ELMO1 as the downstream target for CXCL9-CXCR3, CXCL10-CXCR3, and CXCL11-CXCR3 pairs, CAMK4 as the downstream target for the IFNG-IFNGR1 and IFNG-IFNGR2 pairs, and PCK1 as the downstream target for the IL6-IL6R pair. In addition, CALM1/CALM2/CALM3 were identified as upstream targets for the IL6-IL6R pair.

**Table 3.** KEGG pathways of the ligand (IPF-DEgenes)-receptor pairs and their signaling molecules (IPF-features)

| Pathway | Ligand (IPF-DEgene) | Receptor | Latent disease-omics Feature (IPF-feature) | Position |
| --- | --- | --- | --- | --- |
| hsa04062 | CXCL9 | CXCR3 | ELMO1 | downstream |
| hsa04062 | CXCL10 | CXCR3 | ELMO1 | downstream |
| hsa04062 | CXCL11 | CXCR3 | ELMO1 | downstream |
| hsa04380 | IFNG | IFNGR1 | CAMK4 | downstream |
| hsa04380 | IFNG | IFNGR2 | CAMK4 | downstream |
| hsa04151 | IL6 | IL6R | PCK1 | downstream |
| hsa05163 | IL6 | IL6R | CALM1 | upstream |
| hsa05163 | IL6 | IL6R | CALM2 | upstream |
| hsa05163 | IL6 | IL6R | CALM3 | upstream |

The Pathways are from the KEGG pathways (human). The position indicates whether the corresponding latent disease-omics feature is the downstream or the upstream of the ligand-receptor pair in the corresponding KEGG pathway.

For the 17 ligand (non-lung) – receptor (lung) pairs, we identified four targets (Table 4), ELMO1 as the downstream target for the PF4V1-CXCR3, CXCL9-CXCR3, CXCL10-CXCR3, CXCL11-CXCR3, CXCL13-CXCR3, CXCL13-CXCR5, PF4-CXCR3, and CXCL16-CXCR6 pairs,

**Table 4.**
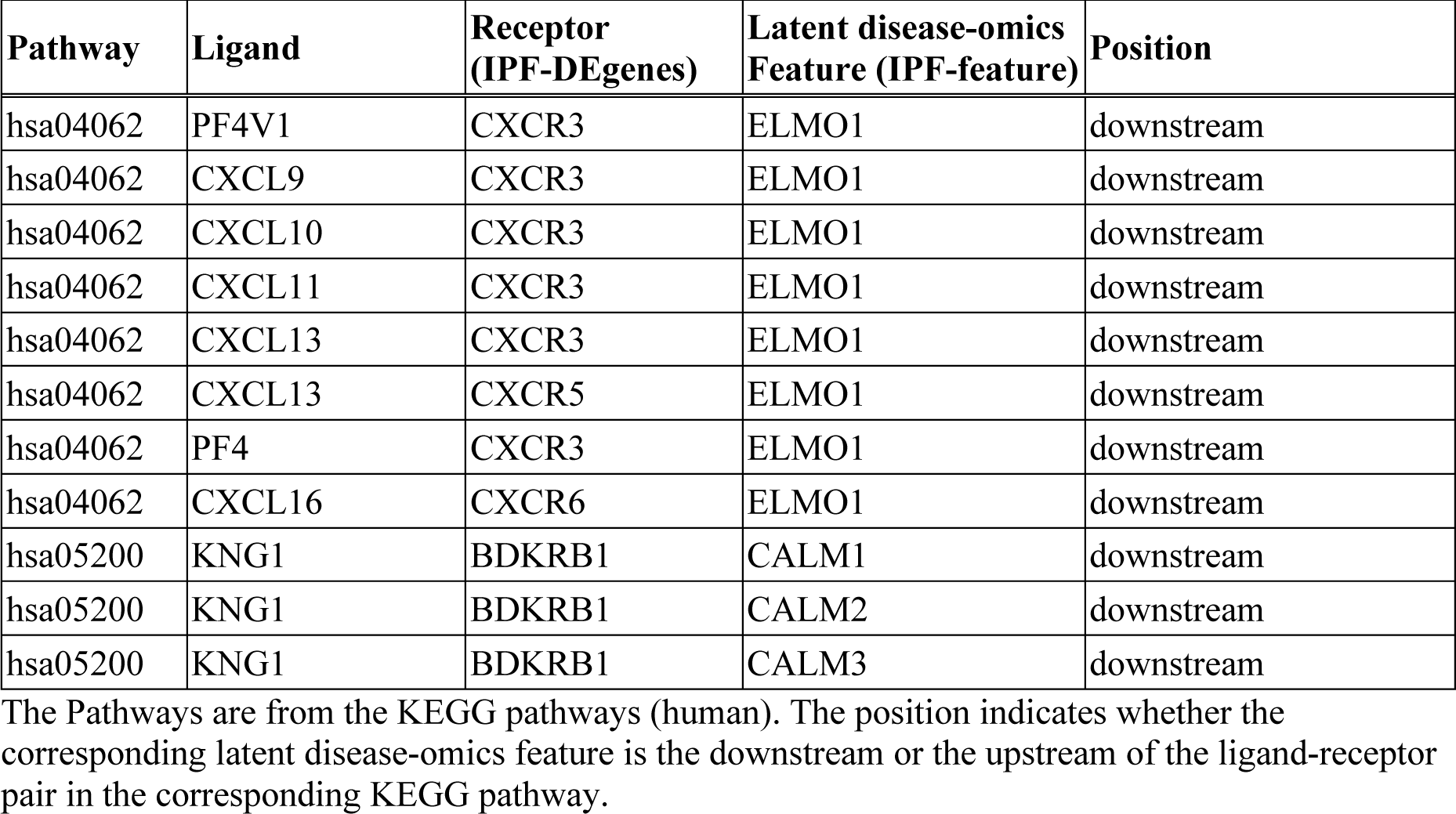
KEGG pathways of the ligand-receptor (IPF-DEgenes) pairs and their signaling molecules (IPF-features)

| Pathway | Ligand | Receptor (IPF-DEgenes) | Latent disease-omics Feature (IPF-feature) | Position |
| --- | --- | --- | --- | --- |
| hsa04062 | PF4V1 | CXCR3 | ELMO1 | downstream |
| hsa04062 | CXCL9 | CXCR3 | ELMO1 | downstream |
| hsa04062 | CXCL10 | CXCR3 | ELMO1 | downstream |
| hsa04062 | CXCL11 | CXCR3 | ELMO1 | downstream |
| hsa04062 | CXCL13 | CXCR3 | ELMO1 | downstream |
| hsa04062 | CXCL13 | CXCR5 | ELMO1 | downstream |
| hsa04062 | PF4 | CXCR3 | ELMO1 | downstream |
| hsa04062 | CXCL16 | CXCR6 | ELMO1 | downstream |
| hsa05200 | KNG1 | BDKRB1 | CALM1 | downstream |
| hsa05200 | KNG1 | BDKRB1 | CALM2 | downstream |
| hsa05200 | KNG1 | BDKRB1 | CALM3 | downstream |

The Pathways are from the KEGG pathways (human). The position indicates whether the corresponding latent disease-omics feature is the downstream or the upstream of the ligand-receptor pair in the corresponding KEGG pathway.

CALM1/CALM2/CALM3 as the downstream target for the KNG1-BDKRB1 pair.

Our aim is to identify the inter-organ mechanism of IPF. The multi-modal generative topic-model found a possible involvement of the liver in this mechanism (Fig. 2). Hence, the ligand-receptor pair(s) that bridge the lung (the primary organ of IPF pathology) and the liver could be such a mechanism. Furthermore, to fulfill this mechanism, the expression of the non-lung component of the ligand-receptor pair should be enriched in the liver.

On the basis of this rationale, we examined the expression patterns of the non-lung components of the ligand-receptor pairs in the liver (Fig. 5, Table S5). The analysis of the multi-organ human scRNA-seq database, *Tabula Sapiens*, identified two pairs, KNG1 (the liver) – BDKRB1 (the lung) and IL6 (the lung) – IL6R/IL6ST (the liver), that could establish the lung-liver inter-organ mechanism. The expression of KNG1, the ligand for BDKRB1, is most enriched in hepatocytes, with lesser expression in the endothelial cells, fibroblasts, intrahepatic cholangiocytes, and T cells. The expression of IL6R/IL6ST, the receptor complex for IL6, is enriched in the endothelial cells of the hepatic sinusoid, intrahepatic cholangiocytes, and hepatocytes.

**Fig. 5.**
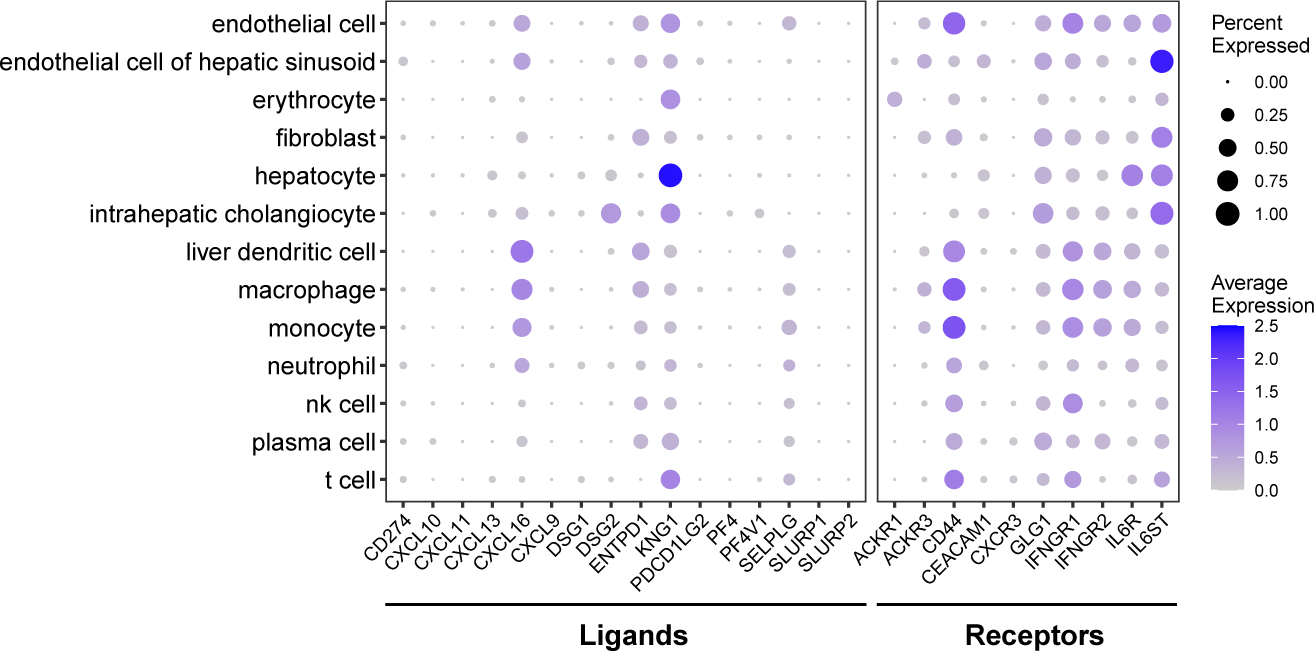
The hepatic expression of the ligands and receptors for the IPF pulmonary receptors and ligands. The level of each ligand and receptor in each cell-type in the liver is shown as dot. The size and the heat-intensity represent the ratio of cells expressing the gene in each cell-type cluster and the mean expression level of log-transformed counts (i.e., log(1 + count per 10,000)), respectively, as shown on the right side of the panel. The raw data are available as Table S5.

Next, we examined the expression patterns of their partner components in the lung (Fig. 6, Tables S6&S7), which were originally found in the 112 IPF-lung DEgenes. The scRNA-seq analysis of IL6, the ligand for the IL6R/IL6ST receptor complex, in the lung using the *Tabula Sapiens* was performed. The result shows that the enrichment of IL6 expression in adventitial cells, endothelial cells, fibroblasts, mesothelial cells, respiratory mucous cells, and smooth muscle cells (Fig. 6A, Table S6). We also examined whether the expression pattern of IL6 in the lung is altered in the IPF patients (Fig. 6B, Table S7). The result identified over 2-fold downregulation of the IL6 expression in endothelial cells and dendritic cells in the IPF lung. In addition, in the lung macrophage, its nearly 2-fold downregulation was also found. While the expression of BDKRB1 is detected more ubiquitously in the lung (Fig. 6A, Table S6), it is nearly 1,000-fold upregulated in the macrophages of the IPF lung, as compared to those of the healthy lung (Fig. 6B, Table S7).

**Fig. 6.**
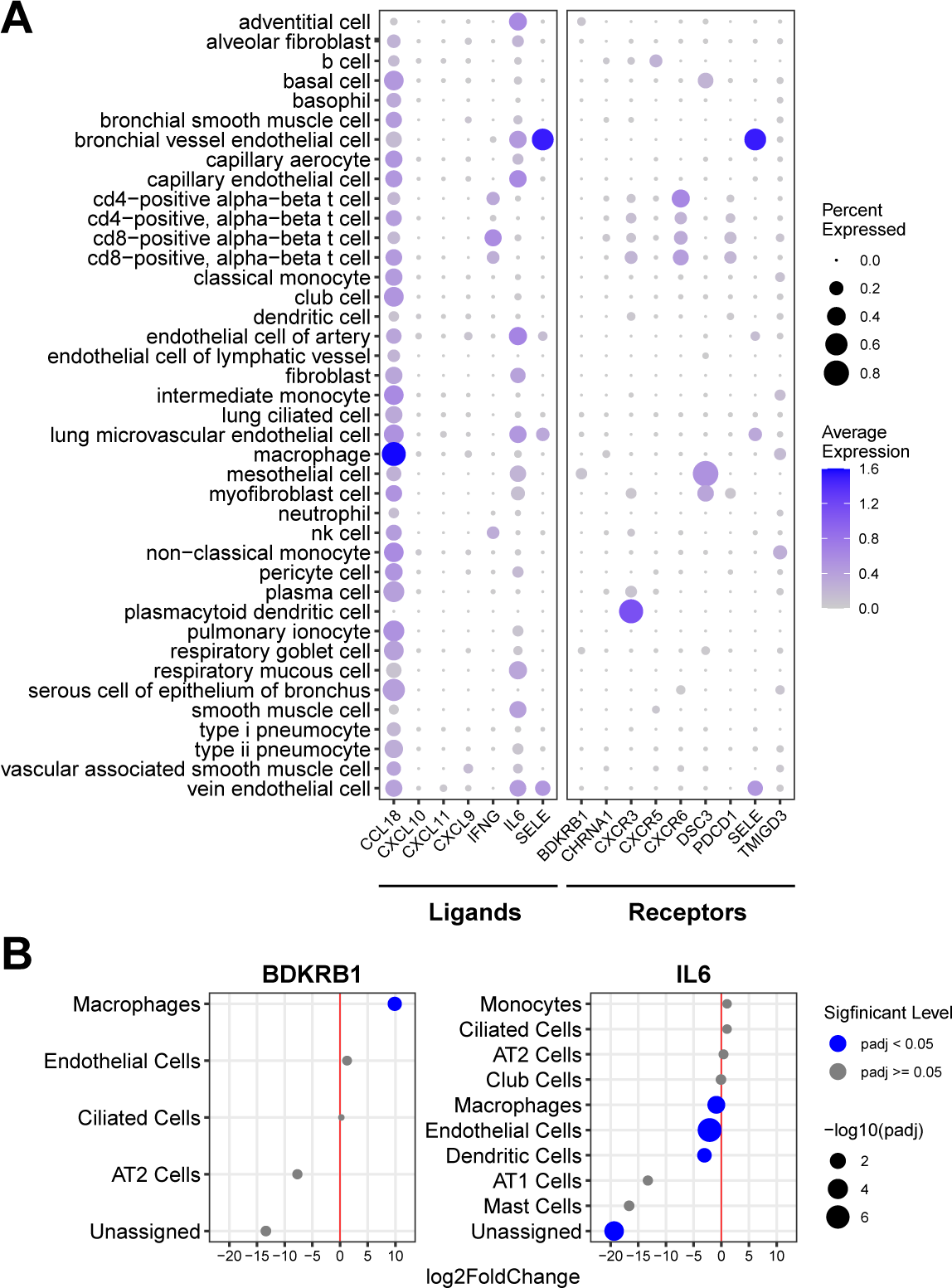
The expression of IL6 and BDKRB1 in the lung. **A:** The level of each ligand and receptor (including IL6 and BDKRB1) in each cell-type in the lung of the healthy subjects (*Tabula Sapiens*) is shown as dot. The size and the heat-intensity represent the ratio of cells expressing the gene in each cell-type cluster and the mean expression level of log-transformed counts (i.e., log(1 + count per 10,000)), respectively, as shown on the right side of the panel. The raw data are available as Table S6. **B:** The differential expression of IL6 and BDKRB1 in each cell-type in the IPF-lung is shown as dot. The cell-types are indicated on the left. The differential expression of IPF vs. non-IPF is indicated as log_2_fold change (“log2FoldChange”). The dot size indicates the statistical significance of the differential expression as −log_10_p-adj (“-log10padj”) – the larger size indicating more significant (i.e., less padj values). The blue and gray colors indicate padj<0.05 and padj≥0.05, respectively. The raw data are available as Table S7.

We also examined the expression patterns of their downstream and upstream signaling targets (Fig. 7). The most significant upregulation of CALM1/CALM2/CALM3, the downstream targets of the KNG1-BDKRB1 signaling and the upstream targets of the IL6-IL6R/IL6ST signaling, was detected in macrophages in the IPF lung (Fig. 7A, Table S6). Lesser but statistically significant upregulation for one or more of these targets was also found in fibroblasts, dendritic cells, T/NKT cells, ciliated cells, monocytes, mast cells, and AT2 cells. Small but statistically significant downregulation was observed for CALM1 in AT1 cells, AT2 cells, and club cells. Such downregulation was also detected for CALM3 in monocytes and dendritic cells.

**Fig. 7.**
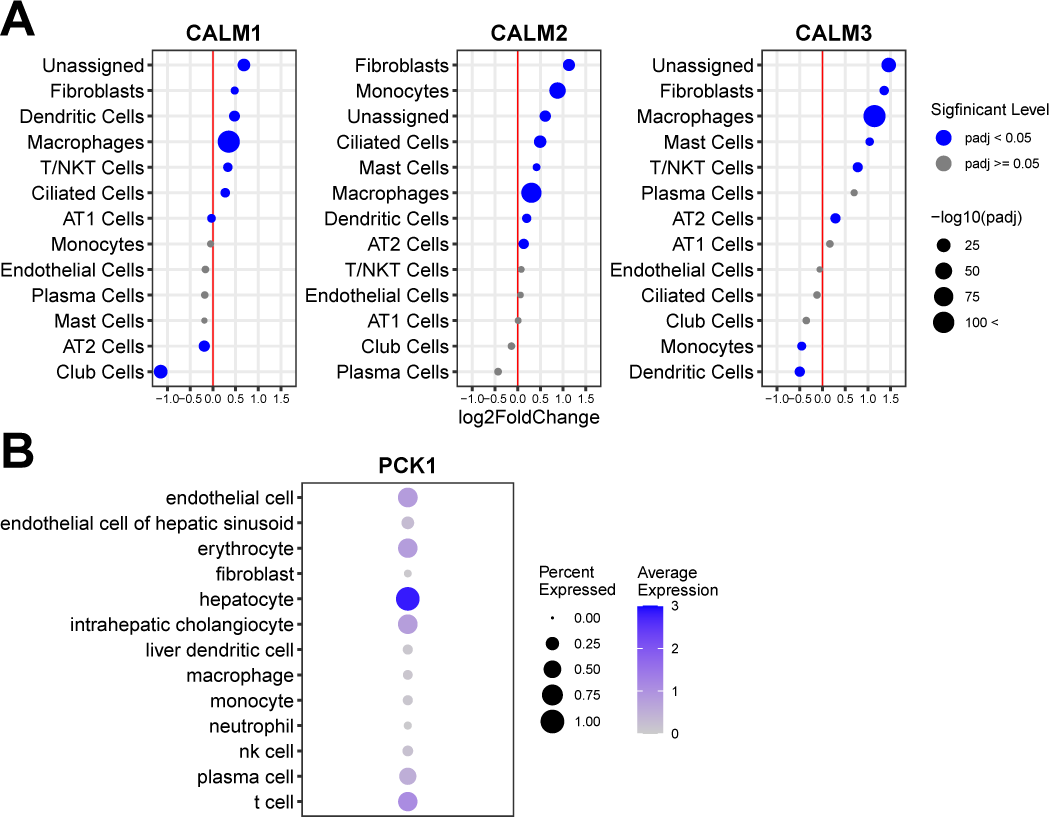
The expression of the signaling targets in the liver and the lung. **A:** The differential expression of CALM1/CALM2/CALM3 in each cell-type in the IPF-lung is shown as dot. The cell-types are indicated on the left. The differential expression of IPF vs. non-IPF is indicated as log_2_fold change (“log2FoldChange”). The dot size indicates the statistical significance of the differential expression as −log_10_p-adj (“-log10padj”) – the larger size indicating more significant (i.e., less padj values). The blue and gray colors indicate padj<0.05 and padj≥0.05, respectively. The raw data are available as Table S7. **B:** The level of PCK1 in each cell-type in the liver is shown as dot. The size and the heat-intensity represent the ratio of cells expressing the gene in each cell-type cluster and the mean expression level of log-transformed counts (i.e., log(1 + count per 10,000)), respectively, as shown on the right side of the panel. The raw data are available as Table S5.

The expression pattern of PCK1, the downstream target of IL6-IL6R/IL6ST signaling, was examined in the liver (Fig. 7B, Table S6). The result shows its highest expression in hepatocytes. Its less abundant expression is detected in endothelial cells, erythrocytes, intrahepatic cholangiocytes, plasma cells, and T cells.

To put these results together, a landscape representing an inter-organ mechanism of IPF emerges (Fig. 8). The logic is as follows: KNG1, expressed in the hepatic cells (Fig. 5), is the systemic ligand for its receptor, BDKRB1 (Table 2). BDKRB1 is also one of the 112 IPF-DEgenes expressed in the pulmonary cells (Table 1, Fig. 6B, Table S1). Hence, the hepatic KNG1 directly interacts with pulmonary BDKRB1 across these organs (Fig. 8). CALM1/CALM2/CALM3, the latent IPF-features (Table S1) are the known downstream targets of KNG1 (ligand) - BDKRB1 (receptor) signaling (KEGG: hsa05200) (Table 4). In addition, CALM1/CALM2/CALM3 are also known upstream signaling components of the IL6 signaling (KEGG: hsa05163) (Table 3). CALM1/CALM2/CALM3 are expressed in the pulmonary cells and their expression is upregulated in the pulmonary macrophages and fibroblasts, etc. of the IPF lung (Fig. 7A). IL6 is one of the IPF-DEgenes (Table S4, Fig. 6B) and is a systemic ligand for its receptor, IL6R/IL6ST (Table 1). IL6R/IL6ST complex is expressed in hepatic cells (Fig. 5).

**Fig. 8.**
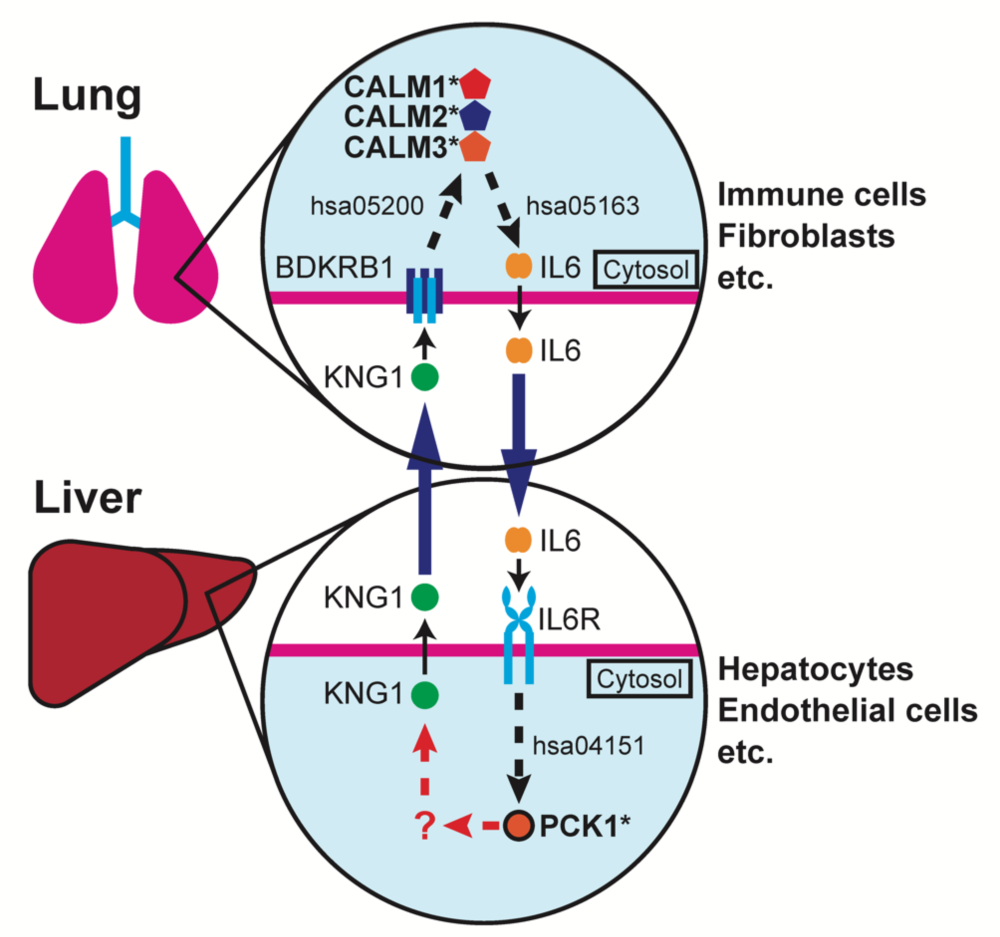
The predicted inter-organ mechanism of IPF. The solid arrows indicate the direct ligand-receptor interactions. The pathway connection (edge) is shown as dashed-arrows indicating the presence of one or more nodes (proteins: ligands, receptors, signaling targets) in between. The corresponding KEGG human pathways for each edge are indicated as hsa numbers. “?” indicates the lack of KEGG pathway connecting the nodes.

Hence, the signal from the liver is transduced to the lung via KNG1 (ligand) – BDKRB1 (receptor) interaction across these organs via CALM1/CALM2/CALM3 to the IL6 signal in the lung (Fig. 8). This pulmonary IL6 signal is transduced back to the liver via the IL6 (ligand) – IL6R/IL6ST (receptor) interaction in the liver (Fig. 8). PCK1, one of the IPF disease-omics features, is a known signaling molecule for the IL6 (ligand) – IL6R/IL6ST (receptor) interaction (KEGG: hsa04151), and it is expressed in the hepatic cells (Fig. 7). Hence the IL6 signal from the IPF-lung is transduced in the liver via PCK1 signaling molecule (Fig. 8). With this logic, the mechanism described in Fig. 8 is generated. In this mechanism, the liver-derived KNG1 activates the CALM1/CALM2/CALM3 signaling pathway via BDKRB1 in the lung. This signal amplifies the expression and/or secretion of IL6 from the lung. The systemic IL6 activates the PCK1 signaling pathway via IL6R/IL6ST in the liver. This feedforward and feedback mechanism across the liver and the lung triggers and/or exacerbates IPF pathogenesis.

## Discussion

### Pre-clinical and clinical evidence

While the results shown in this study are computational, there are mounting pre-clinical and clinical evidence supporting our findings. They are as follows (1&2 below):

1. The latent relatedness of IPF to non-pulmonary diseases

By applying the multi-modal generative topic modeling to the multi-modal disease-omics data of 6,955 human diseases, we identified molecular and genetic relatedness of IPF to non-respiratory diseases such as various types of neoplasm, autoimmune disorders, diabetes, Alzheimer’s disease, rheumatoid arthritis, obesity, cardiovascular diseases (atherosclerosis, arteriosclerosis, hypertensive disease, etc.), systemic lupus erythematosus, and multiple sclerosis (Fig. 3). A possible similarity of IPF to lung cancer is discussed in an editorial article^30^. In this article, RhoGEF mediated epithelial cell transformation (ECT) 2 of AT2 cells in the lung could be a common mechanism between IPF and the lung cancer. Our topic modeling study shows the relatedness of IPF to diverse types of cancer (Fig. 3). As AT2 is a relatively specific resident cell-type in the lung, it is unlikely that the same mechanism is the basis of the relatedness of IPF to other non-lung cancers. However, a possibility of the ECT of other types of epithelial cells in non-lung tissues remains, which could explain the relatedness of IPF to other types of cancer as found in our study.

The relatedness of IPF to diabetes is another finding worth discussion. There are several clinical studies including clinical meta analyses suggesting an association between IPF and diabetes^31–34^. Our computational study also indicates molecular relatedness of IPF to diabetes (Fig. 3). Moreover, other IPF-related diseases found in this study such as Alzheimer’s disease, obesity, cardiovascular diseases, systemic lupus erythematosus are linked to diabetes^35–39^. Furthermore, the two signaling nodes in the inter-organ mechanism of IPF proposed in our study (Fig. 8), CALM1/CALM2/CALM3 (in the lung) and PCK1 (in the liver), are both molecularly linked to diabetes (Table S1). Taken together, IPF may share the same molecular underpinnings with diabetes and other related diseases (i.e., Alzheimer’s disease, obesity, cardiovascular diseases, systemic lupus erythematosus).

2. The inter-organ mechanism of IPF

Our generative computational approach predicts a molecular crosstalk mechanism between the lung and the liver for IPF (Fig. 8). In this mechanism, two secreted systemic factors, KNG1 and IL6, bridge the liver-lung crosstalk. Hence, based on this mechanism, the interference of KNG1-BDKRB1(a receptor for KNG1) and/or IL6-IL6R (a receptor for IL6) interactions could bring a therapeutic benefit to IPF. In this regard, it is worth noting that the blocking IL-6 is shown to attenuate pulmonary fibrosis in mice^40^. In the same study, it is also shown that IPF patients exhibit the increased level of soluble IL6Rα (sIL-6Rα) in their lung tissues. However, the proposed therapeutic mechanism is the blocking of the intra-pulmonary interactions of sIL-6Rα and IL6. Furthermore, possible roles of interleukins in the pathogenesis of pulmonary fibrosis including IPF are recently discussed^41^. In these other studies, the IL6 inhibitory effects in IPF patients are discussed only in the context of heterologous cell-cell crosstalk within the lung tissue. However, as the IL6R is also present in hepatocytes, hepatic endothelial cells, and intrahepatic cholangiocytes (Fig. 5), such IL6-IL6R inhibitory effect could also occur in the liver. Hence, when the therapeutic inhibition of the IL6-IL6R interaction is effective, it is important to consider a possibility that such effects are also through the inhibition of this ligand-receptor interaction outside the lung tissue such as within the liver tissue.

In our inter-organ mechanism, we also propose that the calmodulin pathway (CALM1/CALM2/CALM3) is activated by the liver-derived KNG1 interaction with the lung BDKRB1, which then induces the IL6 pathway (Fig. 8). It is shown that a calmodulin inhibitor, trifluoperazine, exhibits an anti-inflammatory effect in a bleomycin-induced pulmonary fibrosis animal model ^42^. This pre-clinical evidence supports our proposed mechanistic model.

The other signaling node in our inter-organ mechanism is PCK1 (Fig. 8). In our model, PCK1 pathway is activated by the IL6-IL6R interaction in the liver, which is feeds back to the lung pathogenesis of IPF via KNG1-BDKRB1 pathway. Recently, nintedanib, one of the two FDA-approved IPF therapeutics, is shown to attenuate experimental colitis via inhibiting the PCK1 pathway^43^. This study suggests that a part of the therapeutic effect of nintedanib on IPF is via the inhibition of the PCK1 pathway.

### Future perspectives

There are two pending questions in the proposed inter-organ mechanism of IPF. The ligands, receptors, and their signaling targets in this model are co-expressed in multiple cell types in their corresponding organs (Fig. 8). Hence, it remains unknown whether the KNG1-BDKRB1 and IL6-IL6R/IL6ST pathways function within the same cell-type or they interact in *trans* across different cell-types within the same organ.

Another question is whether the IL6-IL6R/IL6ST signal feeds back to KNG1 via PCK1 (Fig. 8). While the signaling of IL6-IL6R/IL6ST to PCK1 is established (hsa04151 KEGG pathway in human), the link of PCK1 to KNG1 remains unknown. Upon the experimental validation of this link, the model becomes a closed feedforward and feedback “loop” across the liver and the lung.

These questions remain for the future studies and their results provide more detailed mechanistic description of IPF. Furthermore, they facilitate the designing of first-in-class therapeutic and/or diagnostic strategies for IPF.

### Conclusions

In this study, we exploited a growing body of multi-modal disease-omics data and a generative computational power to predict an inter-organ mechanism of IPF with the molecular and cellular resolution. Furthermore, our retrospective reference-mining found multiple experimental and clinical evidence in support of the predicted mechanism as described above. Our proposed mechanism is detailed enough, providing a unique opportunity to design hypothesis-driven pre-clinical experiments and/or clinical studies to discover and evaluate first-in-class therapeutic and diagnostic targets for IPF. In addition, our study and results illustrate a computational framework to generate experimentally-testable mechanistic models for other diseases where very little mechanism is known.

## Methods

### Multi-modal generative topic modeling

The multi-modal generative topic modeling approach is as previously described^29^. In this study, we deleted IPF disease-omics data to identify latent IPF-features.

### Organ- and Cell-type expression patterns of the latent IPF-features

The organs and cells where the IPF-features are expressed were identified by organ/cell enrichment analyses using THE HUMAN PROTEIN ATLAS v 21.1.^23–25^, as previously described^29^.

### Latent relatedness of IPF to other diseases

We identified diseases to which IPF is related as previously described^29^. Briefly, the disease-labels of each latent IPF-feature were identified in the Dataset A (Fig. 1) and the frequency of each disease-label was counted. The disease-labels of the higher-frequency are determined as more related to IPF.

### Lung RNA-seq data from patients

The lung tissues were collected from 299 subjects. They consist of 173 idiopathic interstitial pneumonias (IIPs), 76 hypersensitivity pneumonitis (HP), 26 connective tissue diseases (CTD), 24 others (other interstitial lung diseases). The 173 IIPs are further composed of 95 IPF, 41 unclassifiable interstitial pneumonia (UCIP), 28 idiopathic nonspecific interstitial pneumonia (NSIP), 3 idiopathic pleuroparenchymal fibroelastosis (PPFE), and 6 other IIPs. RNA was purified from each sample and processed for RNA sequencing as follows: The lung tissues were sent to TAKARA BIO INC. (Shiga, Japan) for sequencing. At TAKARA BIO INC., a total RNA was purified using NucleoSpin®RNA according to the provided protocol. Before RNA-sequencing, the total RNA for each specimen was checked for quality using Agilent 2000 TapeStation (Agilent Technologies, Santa Clara, CA, USA). The RNA Integrity Numbers (RINs) for all the specimens, as obtained by TapeStation, passed a score of 6.0 or greater. Upon this quality check, mRNA sequencing was performed using the Illumina Sequencer NovaSeq6000 with paired end reads of 150 bps. Read sequences obtained were mapped to genome sequences. Based on the positional information obtained from the mapping and the gene definition file, the gene units were mapped to the genome sequence. The expression level of each gene and transcript was calculated based on the positional information obtained by mapping and the gene definition file and an annotation information was added. The differentially expressed genes (DEgenes) in the lung tissues of IPF vs. all the other pulmonary diseases (UCIP, NSIP, PPFE, other IIPs (labeled as “IIP” in the raw count data), HP, CTD, other interstitial lung diseases (labeled as “Others” in the raw count data) were detected by using an R package, *DESeq2*^29^, with the default parameter settings. The DEgenes were defined as the genes whose absolute values of log2FoldChange are ≥ 1 and also adjusted p-values are < 0.05. These IPF-DEgenes are referred to as “IPF-DEgenes” in this paper.

### Identification of ligand-receptor pairs

From the IPF-DEgenes, those encoding ligand proteins or receptor proteins were identified using a human ligand-receptor combination database, CellChatDB^44^. Furthermore, we identified their corresponding receptor and ligand partners using the same database.

### Generation of an inter-organ map of IPF

The overall design of the approach is described in Fig. 4 (see the Results section for it narrative description). The gene expression analyses in the organs and cells were performed as follows:

The single-cell gene expression across multiple healthy organs and that of the lung tissues of IPF patients were determined using two publicly available human single-cell RNA sequencing (scRNA-seq) datasets: 1) *Tabula Sapiens*, which is a database of multiple healthy tissues^22^, and 2) GSE122960, which is a scRNA-seq dataset of the lung tissues of IPF patients/healthy donors^45^. For the *Tabula Sapiens*, the raw count data and the cell type annotation table were extracted using a Python package, *scanpy*^46^ from the h5ad-formatted data at their FigShare deposit^47^. For the GSE122960, the raw count data were obtained from the GEO deposit and the cell type annotation table was generated using publicly available R program codes^45^. The DEgenes in each cell type of the lung derived from the IPF patients were determined by comparing the scRNA-seq data of the lung tissues derived from the IPF patients and the healthy-donors by *DESeq2*^48^ following the developers’ recommendations for single-cell analysis^49^. Briefly, we first set size-factors by ‘computeSumFactors()’ in the *scran* package^50^. The DESeq was performed by using the likelihood ratio test as significance testing, where we set the ‘DESeq()’ arguments to the following values: test = ‘LRT’, useT = TRUE, minmu = 1e-6, minReplicateForReplace = Inf. The genes were evaluated by the statistical significance level at 0.05 in adjusted p-value.

The KEGG-mining (steps 5&6 in the flow) was conducted as follows:

The KEGG-mining was performed to identify downstream and upstream targets of the ligand-receptor pairs and to determine which of the targets are the IPF-features. First, the gene symbols were converted into KEGG IDs, by first to Entrez IDs using the R function ‘bitr()’ of the R package *clusterProfiler* ^51^, and then to KEGG IDs from the Entrez IDs using KEGG API^52^. Next the KEGG pathways containing these KEGG IDs were extracted and their directed graphs were constructed using KGML^53^. In the graphs, each node was the attribute ‘name’ of the tag <entry>, and each edge started at the node corresponding to the attribute ‘entry1’ of the tag <relation> and ended at the node corresponding to the attribute ‘entry2’ of the same tag <relation>. Using these graphs, we identified the direct and indirect connections between the ligands/receptor and the latent IPF-features.

## Declarations

### Ethics approval and consent to participate

The studies with human subjects and data were approved by the Institutional Review Board of Advanced Telecommunications Research Institute International on behalf of Karydo TherapeutiX, Inc. (Approved Number: HK2101-2101, HK2101-2103, HK2101-2202) and of National Institutes of Biomedical Innovation, Health and Nutrition (Approved Number: 187) and of Kanagawa Cardiovascular and Respiratory Center (Approved Number: KCRC-19-0015).

### Consent for publication

Not applicable

### Availability of data and materials

All raw data of the results are available as supplementary tables in Additional Information 1 of this paper and they are referenced within the manuscript accordingly. The raw count data of the RNA-seq are available at: https://github.com/skozawa170301ktx/IPF The code reported in this paper is available at: https://github.com/skozawa170301ktx/MultiModalDiseaseModeling

### Authors’ contributions

T.N.S. conceived the idea of the project, designed the study, and supervised the overall research project. S.K. designed and performed the multi-modal generative topic modeling analyses. K.T. performed the CellChat DB analyses and gene expression analysis. S.T. performed the KEGG-mining. Y.N.-K., T.O., H.K., T.N. designed the clinical sampling protocol, T.O., H.K., T.N. performed the clinical sampling, Y.N.-K., conducted RNA purification and sequencing, M.N.-I., M.K. performed RNA-seq data preparation. T.N.S., S.K., K.T., S.T., Y.N.-K. wrote the manuscript. All authors reviewed manuscript and approved the final manuscript.

## Acknowledgements

T.N.S., S.K., K.T., and S.T. thank K. Sugisaka, R. Takahashi, R. Ishikawa for their administrative assistance. T.N.S., S.K., K.T., and S.T. are also grateful to the members of Karydo TherapeutiX, Inc. for their supports, advice, and discussion throughout the course of this work. Y.N.-K., T.O., T.N., H.K., M.N.-I., and M.K. thank Yoichi Kurebayashi (Kobe University, Kobe, Japan), Naonori Ueda (RIKEN AIP, Kyoto, Japan), Masanori Shindo (NIBIOHN, Osaka, Japan), Yoshinori Nakamura (NIBIOHN, Osaka, Japan), Hiromitsu Kageyama (NIBIOHN, Osaka, Japan), Michiyo Kawai (NIBIOHN, Osaka, Japan), Akiko Fukagawa (NIBIOHN, Osaka, Japan), Hideyo Kamada (NIBIOHN, Osaka, Japan) for supporting PRISM project.

