## Supplementary Information for "Latent inter-organ mechanism of idiopathic pulmonary fibrosis unveiled by a generative computational approach"

**Supplemental information**

**Supplemental figure**

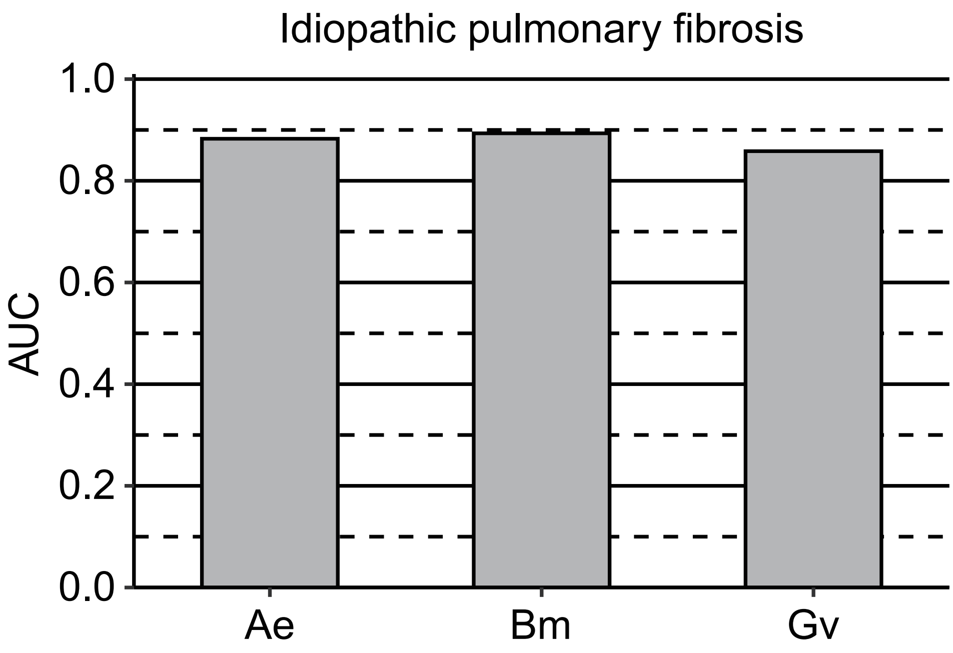


**Fig. S1** The AUC scores of each modality for IPF. The AUC scores are calculated for AlteredExpression (Ae), Biomarker (Bm), GeneticVariation (Gv) modalities for IPF and shown as bar graphs. The bars are shown as the AUC scores of 10x leave-one-modality-out cross-validation.

**Supplementary tables**

**Table S1.** The list of the 83 latent IPF-features.

**Table S2.** The organ and cell enrichment of the 83 IPF-features. The raw data for Fig. 2.

**Table S3.** The disease counts for the 83 IPF-features. The raw data for Fig. 3.

**Table S4.** Raw data of the DESeq2 analysis. The raw data for the 112 IPF-DEgenes in Fig. 4 are included.

**Table S5.** The expression of the ligands, the receptors, and PCK1 in each hepatic cell-type (*Tabula Sapiens*). The raw data for Figs. 5&7B.

**Table S6.** The expression levels of the ligands and receptors in each cell-type in the healthy-lung. The raw data for Fig. 6A.

**Table S7.** The expression levels of IL6, BDKRB1, and CALM1/CALM2/CALM3 in each cell-type in the IPF-lung. The raw data for Figs. 6B&7A.

**List of abbreviations**

***ACKR1:*** Atypical chemokine receptor 1

***ACKR3:*** Atypical chemokine receptor 3

***Ae:*** AlteredExpression

***AI:*** Artificial intelligence

***AT1 cells:*** Alveolar type I cells

***AT2 cells:*** Alveolar type II cells

***AUC:*** Area under the curve

***BDKRB1:*** Bradykinin receptor B1

***Bm:*** Biomarker

***CALM1:*** Calmodulin 1

***CALM2:*** Calmodulin 2

***CALM3:*** Calmodulin 3

***CAMK4:*** Calcium/calmodulin dependent protein kinase IV

***CCL18:*** C-C motif chemokine ligand 18

***CD44:*** Cluster of differentiation 44

***CD274:*** Cluster of differentiation 274

***CEACAM1:*** CEA cell adhesion molecule 1

***CHRNA1:*** Cholinergic receptor nicotinic alpha 1 subunit

***CTD:*** Connective tissue diseases

***CXCL9:*** C-X-C motif chemokine ligand 9

***CXCL10:*** C-X-C motif chemokine ligand 10

***CXCL11:*** C-X-C motif chemokine ligand 11

***CXCL13:*** C-X-C motif chemokine ligand 13

***CXCL16:*** C-X-C motif chemokine ligand 16

***CXCR3:*** C-X-C motif chemokine receptor 3

***CXCR5:*** C-X-C motif chemokine receptor 5

***CXCR6:*** C-X-C motif chemokine receptor 6

***DEgenes:*** Differentially expressed genes

***DSC3:*** Desmocollin 3

***DSG1:*** Desmoglein 1

***DSG2:*** Desmoglein 2

***ECT:*** Epithelial cell transformation

***ELMO1:*** Engulfment and cell motility 1

***ENTPD1:*** Ectonucleoside triphosphate diphosphohydrolase 1

***FDA:*** Food and Drug Administration

***GEO:*** Gene Expression Omnibus

***GLG1:*** Golgi glycoprotein 1

***Gv:*** GeneticVariation

***HP:*** Hypersensitivity pneumonitis

***hsa:*** *Homo sapiens*

***ID:*** Identification

***IFNG:*** Interferon gamma

***IFNGR1:*** Interferon gamma receptor 1

***IFNGR2:*** Interferon gamma receptor 2

***IIP:*** Idiopathic interstitial pneumonias

***IL6:*** Interleukin 6

***IL6R:*** Interleukin 6 receptor

***IL6Rα:*** Interleukin 6 receptor subunit α

***IL6ST:*** Interleukin 6 cytokine family signal transducer

***IPF:*** Idiopathic pulmonary fibrosis

***KEGG:*** Kyoto Encyclopedia of Genes and Genomes

***KGML:*** KEGG Markup Language

***KNG1:*** Kininogen 1

***mRNA:*** messenger RNA (Ribonucleic acid)

***nk cell:*** Natural Killer cell

***NSIP:*** Nonspecific interstitial pneumonia

***padj:*** Adjusted p-value

***PCK1:*** Phosphoenolpyruvate carboxykinase 1

***PDCD1:*** Programmed cell death 1

***PDCD1LG2:*** Programmed cell death 1 ligand 2

***PF4:*** Platelet factor 4

***PF4V1:*** Platelet factor 4 variant 1

***PMC:*** PubMed Central

***PMID:*** PubMed ID

***PPFE:*** Pleuroparenchymal fibroelastosis

***RhoGEF:*** Rho (Rhodopsin) guanine nucleotide exchange factors

***RINs:*** RNA integrity Numbers

***RNA:*** Ribonucleic acid

***RNA-seq:*** RNA sequencing

***scRNA-seq:*** single-cell RNA sequencing

***SELE:*** Selectin E

***SELPLG:*** Selectin P ligand

***sIL-6Rα:*** Soluble interleukin 6 receptor subunit α

***SLURP1:*** Secreted LY6/PLAUR domain containing 1

***SLURP2:*** Secreted LY6/PLAUR domain containing 2

***TGF-beta:*** Transforming growth factor-beta

***T/NKT cells:*** T/Natural Killer T cells

***TMIGD3:*** Transmembrane and immunoglobulin domain containing 3

***UCIP:*** Unclassifiable interstitial pneumonia
